# Murine *Surf4* is essential for early embryonic development

**DOI:** 10.1101/541995

**Authors:** Brian T. Emmer, Paul J. Lascuna, Emilee N. Kotnik, Thomas L. Saunders, Rami Khoriaty, David Ginsburg

## Abstract

Newly synthesized proteins co-translationally inserted into the endoplasmic reticulum (ER) lumen may be recruited into anterograde transport vesicles by their association with specific cargo receptors. We recently identified a role for the cargo receptor SURF4 in facilitating the secretion of PCSK9 in cultured cells. To examine the function of SURF4 *in vivo*, we used CRISPR/Cas9-mediated gene editing to generate mice with germline loss-of-function mutations in *Surf4*. *Surf4*^*+/-*^ mice exhibited grossly normal appearance, behavior, body weight, fecundity, and organ development and demonstrated no significant alterations in circulating plasma levels of PCSK9, apolipoprotein B, or total cholesterol. *Surf4*^*-/-*^ mice exhibit embryonic lethality, with complete loss of all *Surf4*^*-/-*^ offspring between embryonic days 3.5 and 9.5. Taken together with the much milder phenotypes of PCSK9 or apolipoprotein B deficiency in mice, these findings imply the existence of additional SURF4 cargoes or functions that are essential for murine early embryonic development.

## INTRODUCTION

The coatomer protein complex II (COPII) coat assembles on the cytoplasmic surface of ER exit sites to drive the formation of membrane-bound transport vesicles. Efficient recruitment of proteins and lipids into these vesicles occurs via physical interaction with the COPII coat^1^. For cargoes accessible on the cytoplasmic surface of the ER membrane, this interaction may be direct. For soluble cargoes in the ER lumen, however, transmembrane cargo receptors serve as intermediaries for this interaction^2^. Although thousands of human proteins traffic through the secretory pathway, a corresponding cargo receptor has been identified for only a few, and the size and identity of the cargo repertoire for each individual cargo receptor remains largely unknown.

Through unbiased genome-scale CRISPR screening, we recently discovered a role for the ER cargo receptor SURF4 in the secretion of PCSK9, a protein that modulates mammalian cholesterol homeostasis through its negative regulation of the LDL receptor^3^. Consistent with a role as a PCSK9 cargo receptor, SURF4 was found to localize to the ER and ERGIC compartments, to physically associate with PCSK9, and to promote the ER exit and extracellular secretion of PCSK9. These experiments relied on heterologous expression of PCSK9 in HEK293T cells, however, and the physiologic relevance of this interaction *in vivo* remains uncertain. Additionally, although SURF4 deletion did not affect the secretion of a control protein, alpha-1 antitrypsin, a broader role for SURF4 in protein secretion remains possible and is supported by the recent identification of other potential cargoes including apolipoprotein B, growth hormone, DSPP, and amelogenin^4,5^.

To investigate the physiologic functions of SURF4, we generated mice with targeted disruption of the *Surf4* gene. We found that partial loss of SURF4 in heterozygous mice caused no significant changes to circulating levels of PCSK9, apolipoprotein B levels, or total cholesterol, but that complete genetic deletion of *Surf4* resulted in early embryonic lethality.

## RESULTS

### Generation of mice with germline deletion of *Surf4*

The *Mus musculus Surf4* gene is composed of 6 exons and 5 introns spanning approximately 14 kb in the tightly clustered surfeit gene locus on chromosome 2^6,7^. We targeted exon 2 of *Surf4* for CRISPR/Cas-mediated mutagenesis (Figure 1A), verified sgRNA efficiency in embryonic stem (ES) cells (Figure 1B-C), and generated mice from microinjected zygotes. Sanger sequencing identified 4 of 57 mice with disruption of the target site (Figure 1D). These mice were then mated to C57BL/6J wild-type mice and their progeny genotyped, confirming germline transmission for each of the 4 alleles (Figure 1E). Of these, 2 alleles introduced a frameshift deletion leading to an early termination codon, with the other alleles containing in-frame deletions of 3 and 6 DNA base pairs, respectively.

**Figure 1:**
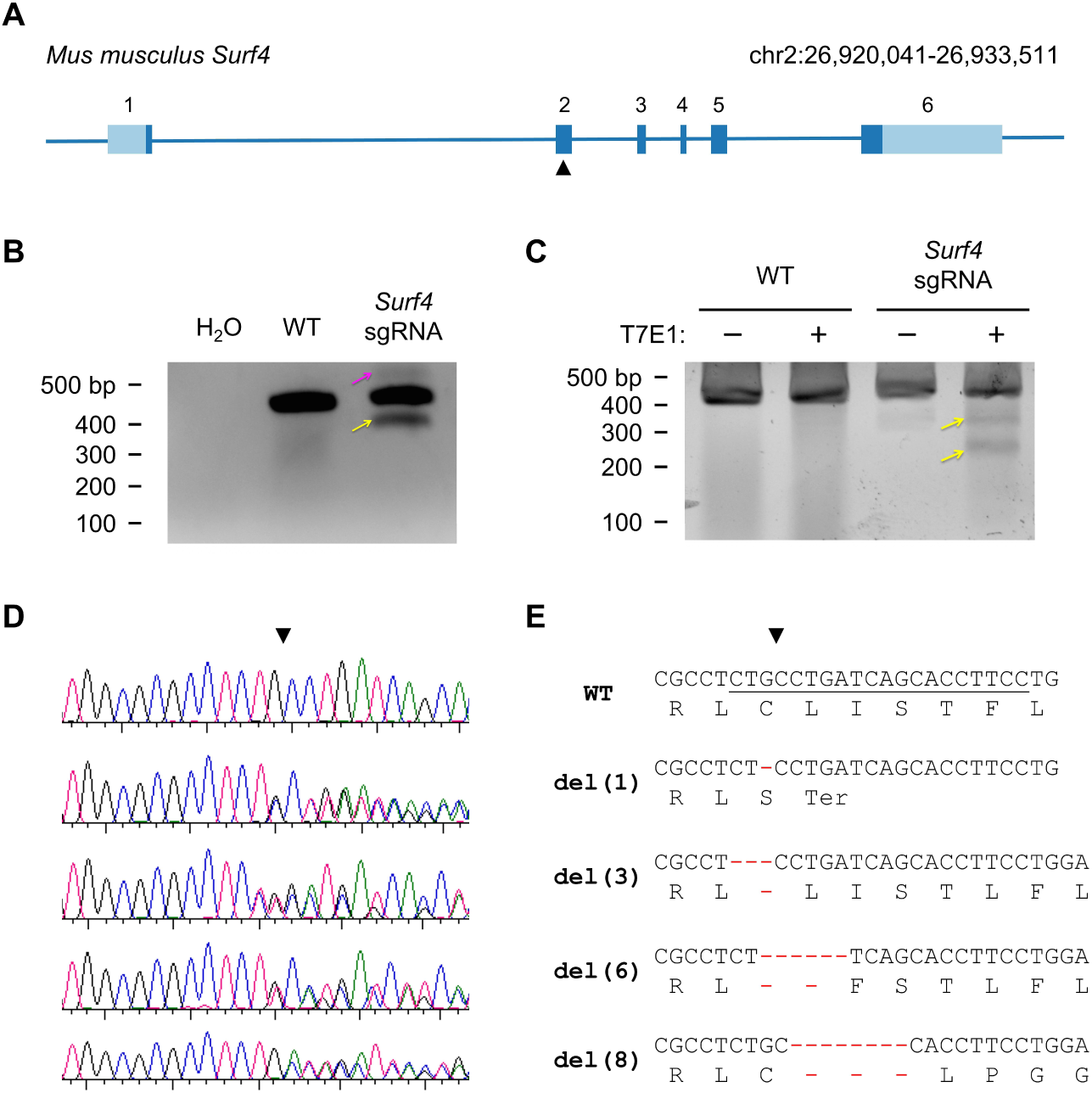
Generation of *Surf4* mutant alleles. (A) *Surf4* gene structure. Exons are shaded light blue for UTRs or dark blue for coding sequence. The target site for the sgRNA used for oocyte editing is indicated by the black triangle. (B) Mouse ES cells were either untreated or electroporated with plasmids for CRISPR/Cas9 disruption of the *Surf4* target site. PCR amplification of genomic DNA or water control across the *Surf4* target site revealed higher and lower molecular weight DNA fragments suggestive of nonhomologous endjoining repair of Surf4 indels. (C) The major PCR product was gel purified and subjected to T7 endonuclease I digestion. T7E1 digestion produced novel DNA fragments (arrows) indicating the presence of insertions/deletions in *Surf4* exon 2. Wild type DNA was resistant to T7E1 digestion. (D) Sanger sequencing chromatograms of *Surf4* target site amplicons of progeny from matings between *Surf4*-targeted founder mice and wild-type C57BL6/J mice. (E) DNA and predicted protein sequences of each individual allele generated by CRISPR/Cas9 gene-editing of *Surf4*.

### Effect of SURF4 haploinsufficiency on cholesterol regulation

*Surf4*^+/-^mice were present in expected Mendelian ratios at weaning (Table 1) and exhibited grossly normal appearance, behavior, and organ development by necropsy. To further assess the impact of haploinsufficiency for SURF4, we measured plasma PCSK9, cholesterol, and apolipoprotein B levels in *Surf4*^+/-^mice harboring the frameshift-causing del(1) allele and in *Surf4*^+/+^ litter-mate controls. No significant differences were observed for each of two independent blood draws from each mouse (Figure 2).

**Table 1:**
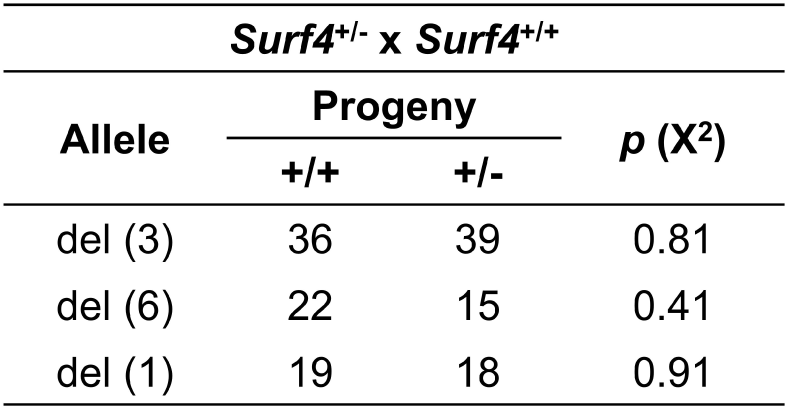
Heterozygosity for *Surf4* mutant alleles is present at expected Mendelian ratios. Mice heterozygous for the indicated *Surf4* allele were crossed with wild-type C57BL/6J mice and the resulting litters genotyped for *Surf4*. The proportion of mice with the heterozygous mutant genotype was compared to expected Mendelian ratios by the chi-square test.

| *Surf4<sup>+/-</sup> x Surf4<sup>+/+</sup>* |  |  |  |
| --- | --- | --- | --- |
| Allele | Progeny |  | *p* (X<sup>2</sup>) |
|  | +/+ | +/- |  |
| del (3) | 36 | 39 | 0.81 |
| del (6) | 22 | 15 | 0.41 |
| del (1) | 19 | 18 | 0.91 |

**Figure 2.**
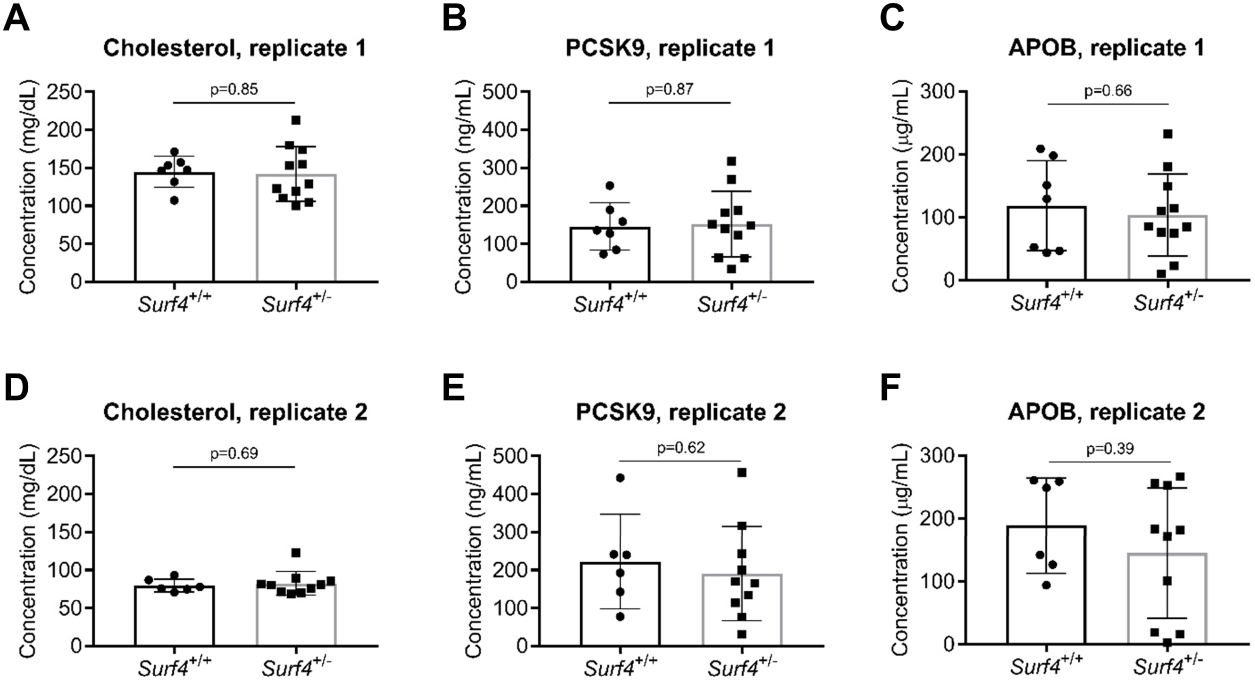
*Surf4* haploinsufficiency does not affect PCSK9, ApoB, LDL receptor, or plasma cholesterol levels when fed a normal or high-fat diet. *Surf4*^*+/-*^ mice harboring the del(1) allele were compared to litter-mate *Surf4*^*+/+*^ control mice for plasma levels on two separate blood draws (A – C and D – F) of total cholesterol (A, D), PCSK9 (B, E), and ApoB (C, F). Significance testing was calculated by Student’s t-test between genotype groups.

### *Surf4* deletion causes early embryonic lethality

The genotypes of pups from intercrosses of *Surf4*^+/-^mice carrying each of 3 independent *Surf4* deletion alleles are shown in Table 1. Though the expected number of *Surf4*^+/+^ and *Surf4*^+/-^pups were observed at the time of weaning, no *Surf4*^-/-^mice were recovered (Table 2). The genotype results from timed matings performed for the del(1) allele are shown in Table 3. No *Surf4*^-/-^embryos were recovered at E9.5 or later. Analysis of E3.5 blastocysts generated by *in vitro* fertilization revealed the expected number of *Surf4*^-/-^genotypes. Thus, homozygous deficiency of *Surf4* results in embryonic lethality occurring sometime between E3.5 and E9.5.

**Table 2:**
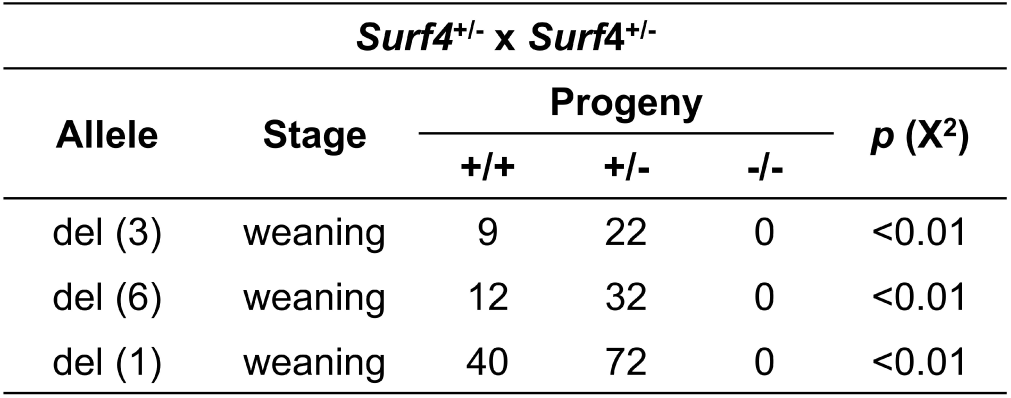
Germline deletion of *Surf4* causes embryonic lethality. Mice heterozygous for the indicated *Surf4* allele were mated and progeny were genotyped for *Surf4* at weaning. The proportion of mice with the homozygous null genotype was compared to expected Mendelian ratios by the chi-square test.

| *Surf4<sup>+/-</sup> x Surf4<sup>+/-</sup>* |  |  |  |  |  |
| --- | --- | --- | --- | --- | --- |
| Allele | Stage | Progeny |  |  | *p* (X<sup>2</sup>) |
|  |  | +/+ | +/- | -/- |  |
| del (3) | weaning | 9 | 22 | 0 | <0.01 |
| del (6) | weaning | 12 | 32 | 0 | <0.01 |
| del (1) | weaning | 40 | 72 | 0 | <0.01 |

**Table 3:** Germline deletion of Surf4 causes lethality between embryonic day 3.5 and 9.5. Timed matings were performed between *Surf4*^+/-^mice harboring a 1 nucleotide deletion and embryos harvested at E9.5 and later. For analysis at E3.5, blastocysts were collected following *in vitro* fertilization of eggs from *Surf4*^+/-^females with sperm from *Surf4*^+/-^males. The proportion of mice with the homozygous null genotype was compared to expected Mendelian ratios by the chi-square test.

| *Surf4<sup>+/-</sup> x Surf4<sup>+/-</sup>* |  |  |  |  |  |
| --- | --- | --- | --- | --- | --- |
| Allele | Stage | Progeny |  |  | *p* (X<sup>2</sup>) |
|  |  | +/+ | +/- | -/- |  |
| del (1) | weaning | 40 | 72 | 0 | <0.01 |
|  | E14.5 | 2 | 3 | 0 | >0.99 |
|  | E12.5 | 3 | 12 | 0 | 0.100 |
|  | E9.5 | 5 | 15 | 0 | 0.047 |
|  | E3.5 | 1 | 7 | 4 | >0.99 |

## DISCUSSION

Identification of the molecular machinery underlying eukaryotic protein secretion has been elucidated by elegant work in model systems including yeast and cultured mammalian cells. Recent characterizations of mice with genetic deficiency of COPII components have extended these findings to mammalian physiology, revealing a variety of complex phenotypes^8-17^. Comparatively little is known about the physiologic role of mammalian cargo receptors *in vivo*. In humans, genetic deletion of either subunit of a cargo receptor complex, LMAN1/MCFD2, results in a rare bleeding disorder due to the impaired secretion of coagulation factors V and VIII^18,19^. Genetic deletion of *Lman1* in mice recapitulates impaired clotting factor secretion of the human disease^20^.

We set out to investigate the physiologic function of murine SURF4 due to its putative roles in the secretion of PCSK9^3^ and apolipoprotein B^5^, both of which play central roles in mammalian cholesterol regulation. In cultured cells, secretion defects of PCSK9^3^ and apoB^5^ are observed upon complete deletion of *SURF4*. By contrast, we found that partial reduction of SURF4 in *Surf*^*+/-*^ mice was associated with normal circulating levels of PCSK9, apolipoprotein B, and total cholesterol. This observation is similar to the unaffected serum levels of LMAN1 cargo proteins in *Lman1*^+/-^mice^20^. Together these findings suggest that under basal conditions, these ER cargo receptors may be present in stoichiometric excess of their cargoes, enabling accommodation of flux in the rates of synthesis and secretion.

Consistent with partial loss of SURF4 being well-tolerated, SURF4 loss-of-function variants have been detected in the Genome Aggregation Database (gnomAD), though at less than expected frequencies (observed/expected ratio 90% confidence interval 0.07-0.55)^21^. Of note, previous human genome-wide association studies for lipid traits failed to detect a significant signal near the SURF4 gene^22^.

Our attempts to generate adult mice with complete loss of *Surf4* were precluded by the embryonic lethality caused by *Surf4* deletion. This phenotype was unlikely to have been caused by a linked spontaneous or off-target CRISPR-generated passenger mutation^23^ as it was observed for each of 3 independent alleles. Timed matings revealed that loss of *Surf4*^-/-^embryos occurs between E3.5 and E9.5. The mechanism for this observation is unclear. Deficiency of the SURF4 homologue SFT-4 is similarly associated with embryonic lethality in *C. elegans*^5^, but SURF4 is not essential for cellular viability in cultured HEK293T cells^3,4^ and its homologue, Erv29p, is dispensable in yeast^24,25^. Deficiencies of PCSK9 or apoliporotein B alone cannot account for this developmental phenotype, given that PCSK9^-/-^mice are viable^26^ and that ApoB^-/-^mice survive past E9.5^27^. Likewise, mice with genetic deletion of other putative SURF4 cargoes growth hormone^28^, amelogenin^29^, and dental sialophosphoprotein^30^ are viable. The embryonic lethality of SURF4 may therefore result from additive effects of disrupted secretion of known cargoes, or the presence of additional unknown SURF4 cargoes or functions that are essential for early embryonic development. A role for SURF4 in global protein secretion is unlikely, as previous studies demonstrated no effect of SURF4 deletion on the secretion of a number of other proteins ^3,4^. A broader role for SURF4 in the secretion of additional unknown cargoes however is suggested by the observation that SURF4 has been evolutionarily conserved in ancient organisms lacking homologues of the putative mammalian cargo, PCSK9. An N-terminal tripeptide “ER-ESCAPE motif”, present on a vast number of potential cargoes, has recently been proposed to mediate cargo recruitment by SURF4^4^. A comprehensive identification of SURF4 cargoes and the nature of their interaction with SURF4 will clarify the function of SURF4 in cholesterol regulation and in mammalian development.

## MATERIALS AND METHODS

### Generation of *Surf4* mutant mice

We used CRISPR/Cas9 technology^31,32^ to generate a new genetically modified mouse strain with a *Surf4* gene knockout. The presence of a premature termination codon in exon 2 is predicted to result in loss of protein expression due to nonsense mediated decay of mRNA^33^. A single guide RNA (sgRNA) target and protospacer adjacent motif was identified in exon 2 (ENSMUSE00000232711.1) with CRISPOR^34^. The sgRNA is 5’ CTGCCTGATCAGCACCTTCC TGG 3’ on the non-coding strand (chromosome 2; coordinates 26926892-26926911) with a predicted cut site 47 bp downstream of the first exon 2 codon. The sgRNA target was cloned into plasmid pX330 (Addgene.org plasmid #42230, a kind gift of Feng Zhang) as described^35^. The sgRNA was validated in mouse JM8.A3 ES cells^36^prior to use for mouse zygote microinjection. The sgRNA plasmid (15 μg) was electroporated into 8 × 10E6 ES cells. To the electroporation 5 μg of a PGK1-puromycin resistance plasmid^37^ was added for transient puromycin selection (2 μg/ml puromycin applied 48-72 hours after electroporation). ES cell culture and electroporation was carried out as described^38^. After selection, DNA was extracted from surviving cells, PCR was used to amplify the sequences across the sgRNA cut site, and T7 endonuclease 1 assays were used to detect small deletions/insertions at the predicted Cas9 DNA cut site^39^. The circular sgRNA plasmid was resuspended in microinjection buffer as described^40^. The plasmid mixture was used for pronuclear microinjection of zygotes obtained from the mating of superovulated C57BL/6J female mice (The Jackson Laboratory Stock No. 0006640) and C57BL/6J male mice as described^41^. A total of 305 zygotes were microinjected, 285 zygotes were transferred to pseudopregnant B6D2F1 female mice (The Jackson Laboratory Stock No. 100006). 18 mouse pups were born and four of them transmitted gene edited *Surf4* alleles.

### Mouse genotyping

*Surf4* genotyping was performed by PCR of genomic DNA with primers mSurf4-ex2-for [TGCTGAGGGCCTCTCTGTCT] and mSurf4-ex2-rev [CAGGTAGCCACAGCTCCAGG]. Sanger sequencing was performed with the same genotyping primers and chromatograms were inspected both manually and by automated deconvolution^42^ to determine the presence or absence of target site indels.

### Analysis of *Surf4*^+/-^mice

Mice were housed and monitored in accordance with University of Michigan Unit of Laboratory Animal Medicine (ULAM) guidelines. Blood was collected at 6-12 weeks of age by retro-orbital bleeding into heparin-coated collection tubes from mice anesthetized with isoflurane. Plasma samples were collected by centrifugation at 2,000 g for 10 min at 4°C. A second blood collection was performed 1 week following the initial collection. Plasma samples were analyzed by total cholesterol colorimetric assay (SB-1010-225, Fisher Scientific, Hampton NH) and ELISAs for PCSK9 (MPC900, R&D Systems, Minneapolis MN) and apolipoprotein B (ab230932, Abcam, Cambridge UK).

### Timed matings and *in vitro* fertilization

For analysis of embryonic day 9.5 and later, *Surf4*^+/-^male and female mice were co-housed overnight and females with copulatory vaginal plugs the following morning were assigned embryonic day 0.5. Pregnant females were then sacrificed at indicated time points and genomic DNA prepared from isolated embryos. For analysis of embryonic day 3.5, *Surf4*^+/-^females were superovulated with anti-inhibin serum as described^43^. The collected oocytes fertilized with sperm from *Surf4*^+/-^males as described^44^. Resulting fertilized eggs were maintained in cell culture in KSOM medium (Zenith Biotech) until harvesting and genomic DNA preparation from blastocysts at embryonic day 3.5.

